# Recording of DNA binding events during gut commensalism reveals the action of a repurposed *Candida albicans* regulatory network

**DOI:** 10.1101/2020.06.24.169920

**Authors:** Jessica N. Witchley, Cedric A. Brimacombe, Nina V. Abon, Suzanne M. Noble

**Affiliations:** Department of Microbiology and Immunology, University of California San Francisco, CA 94143; Department of Medicine, Division of Infectious Diseases, University of California San Francisco, CA 94143

## Abstract

*Candida albicans* is a central fungal component of the human gut microbiota and an opportunistic pathogen. Two *C. albicans* transcription factors, Wor1 and Efg1, control its ability to colonize the mammalian gut. They are also master regulators of an epigenetic switch required for mating. Here, we show that six additional mating regulators influence gut commensalism. Using an adapted Calling Card-Seq protocol to record *Candida* transcription factor DNA binding events in the host, we validated these relationships during murine gut colonization. Finally, by comparing the in-host transcriptomes of regulatory mutants with enhanced vs. diminished commensal fitness, we identified a set of candidate commensalism effectors. These include Cht2, a GPI-linked chitinase whose gene is bound by Wor1, Czf1, and Efg1 *in vivo* and that we show to promote commensalism. We conclude that the network required for a *C. albicans* sexual switch is biochemically active in the host digestive tract, where it is repurposed to direct commensalism.

## INTRODUCTION

*Candida albicans* is the major fungal commensal and opportunistic pathogen of humans. As a pathogen, *C. albicans* is responsible for syndromes ranging from simple vaginitis to highly morbid infections of blood and internal organs. Whereas life-threatening disease is primarily restricted to patients with underlying risk factors such as immunodeficiency, indwelling catheters, and exposure to broad-spectrum antibiotics (1,2), most humans engage in commensal interactions with *C. albicans,* which composes part of normal gut, skin, and genitourinary microbiota. Observations in mice suggest that colonization with *C. albicans* may even benefit the host by promoting immunity to fungal, viral, and bacterial pathogens (3,4). For example, recent work shows that the majority of human anti-fungal Th17 cells, which provide critical mucosal immunity to extracellular pathogens, react with *Candida albicans* (3). Given the diverse contributions of *C. albicans* to human health and disease, there is surprisingly little understanding of the mechanisms that underlie these behaviors.

Our group previously identified two *C. albicans* transcription factors, Efg1 and Wor1, that play opposing roles in the regulation of gut commensalism (**Figure 1A**) (5). Efg1 inhibits commensal fitness, such that an *efg1* null mutant will strongly outcompete wild-type *C. albicans* in mouse models of gut colonization (5–7). In contrast, Wor1 promotes commensalism, such that *wor1* mutants are rapidly outcompeted by wild type (5). Consistent with a special role for Wor1 in the gastrointestinal niche, *WOR1* gene expression—which is barely detectable under *in vitro* conditions—is upregulated ~10,000-fold when cells are propagated in the gut (5). Further, forced overexpression of *WOR1* (*WOR1*^OE^) via a heterologous promoter confers enhanced commensal fitness (5). Interestingly, the fitness advantage of *WOR1*^OE^ cells is not apparent until after 5 to 10 days of host colonization, when they undergo a stochastic but stable transition from standard, oval-shaped “white” morphology to an elongated “GUT” (<u>G</u>astrointestinally Ind<u>U</u>ced <u>T</u>ransition) phenotype (5). Unlike white cells, GUT cells exhibit a distinct metabolism that is oriented towards nutrients available in the mammalian digestive tract. and these cells are immediately hypercompetitive when introduced into naïve animals (5). We hypothesize that wild-type *C. albicans* also undergoes a white-to-GUT switch while within the commensal niche, but reverts to white morphology upon exit from the gut, when signals required for *WOR1* expression are removed.

**Figure 1.**
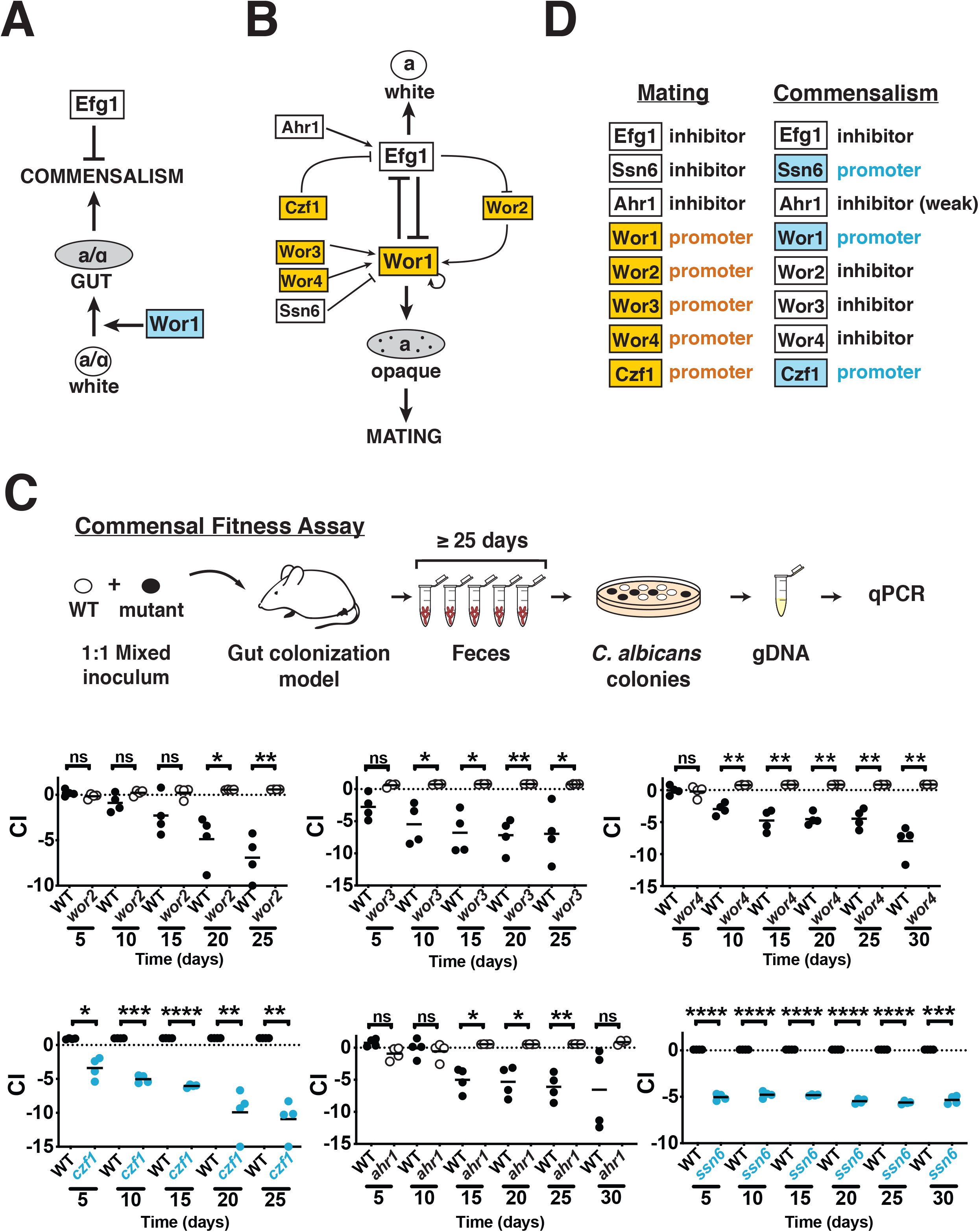
*C. albicans* sexual regulators control commensal fitness. A. Regulation of gut commensalism. Based on analysis of *C. albicans* mutants in a mouse model of gut colonization, Efg1 inhibits (bar) and Wor1 promotes (arrow) fungal commensalism (5). Moreover, within the mammalian digestive tract, Wor1 promotes a white-to-GUT developmental switch that confers enhanced fitness in this niche. B. Regulation of mating. Based on assessment of *C. albicans* mutants for the ability to mate and to switch between white and opaque cell types under *in vitro* conditions, Efg1 inhibits and Wor1 promotes mating and switching. Further, additional transcription factors have been demonstrated to interact with Efg1 and Wor1 in a complex regulatory circuit (8–10,13–15,17,18,46,47). C. Besides Efg1 and Wor1, six additional sexual regulators influence the commensal fitness of *C. albicans. wor2, wor3, wor3, czf1, ahr1,* and *ssn6* mutants were competed against wild type in the murine gut colonization model, and the competitive index (CI) of each strain was monitored 25-30 days. Statistical comparisons were performed using the paired student’s t-test, * p<0.05, ** p<0.01, *** p<0.001, **** p<0.0001. D. Comparison of the roles of eight fungal transcription factors in mating vs. commensalism.

Besides their roles in commensalism, Efg1 and Wor1 are also “master regulators” of *C. albicans* mating (8–11). Wor1 promotes and Efg1 opposes the white-to-“opaque” morphological transition, an epigenetic switch that confers sexual competency in this species. Most *C. albicans* strains are incapable of mating because they express **a**1-α2, a potent transcriptional inhibitor of *WOR1* that is encoded by two heterologous alleles of the Mating Type-Like Locus, *MTL***a** (**a**) and *MTL*α (α). To mate, diploid **a**/α white cells must lose one allele of *MTL,* then switch to the opaque cell type, before opaque **a** and α cells fuse to generate tetraploid **a**/α progeny. In addition to Wor1 and Efg1, other transcription factors have also been discovered to control sexual competency through a complex, self-reinforcing regulatory circuit that has been deduced from genetic epistasis experiments (**Figure 1B**, (12–16)). Further, chromatin immunoprecipitation (ChIP) experiments indicate that Wor1, Efg1, Wor2, Wor3, Wor4, Czf1, Ahr1, and Ssn6 engage in extensive cross-regulation, with each transcription factor binding to its own promoter as well as to those for most or all of the other regulators in the circuit (**Figure S1A** and **S1B**; (9–11,13–19)).

Given that an antagonistic relationship between Wor1 and Efg1 is conserved between mating and commensalism, we hypothesized that additional regulators of mating might help to regulate commensalism. To test this idea, we performed a systematic analysis of *wor2, wor3, wor4, czf1, ahr1,* and *ssn6* null mutants in a murine gut colonization model. Remarkably, all six transcription factors exert significant effects on commensal fitness, although the activities of several regulators differ between sex and commensalism. To validate these relationships in *Candida* colonizing the murine gut, we adapted a transposon-based Calling Card-seq method to record *in vivo* regulatory targets of DNA-binding proteins. Finally, we identified a set of candidate commensalism effectors by comparing the transcriptomes of regulatory mutants with enhanced versus diminished commensal fitness. Among these, we found that *CHT2* (which encodes a GPI-linked fungal chitinase) is a direct target of Wor1, Czf1, and Efg1 and is required for normal fitness in animals. Our work establishes that a transcriptional circuit that determines *C. albicans* mating competency under *in vitro* conditions is repurposed in the host to control its ability to flourish as a gut commensal.

## RESULTS

### Regulators of sexual switching control commensal fitness

To determine whether sexual regulators besides Wor1 and Efg1 play roles in commensalism, we generated null mutants affecting Czf1, Wor2, Wor3, Wor4, Ahr1, and Ssn6 (**Figure 1A**). Homozygous disruptants of each open reading frame (ORF) were created in SN152, the same genetic background used for our previously reported *wor1, efg1,* and *WOR1*^OE^ strains (5). The competitive fitness of each mutant was determined using the mouse gastrointestinal colonization model depicted in **Figure 1C**. Immunocompetent BALB/c mice were treated with broad spectrum antibiotics (ampicillin and gentamicin in drinking water) for one week, followed by gavage with a 1:1 mixture of wild-type *C. albicans* (WT) and a transcription factor mutant. Antibiotics were continued, and the relative abundance of fungal strains recovered from host feces over ≥25 days was determined by qPCR. The competitive index (CI) of each mutant was calculated as the log_2_ ratio of its relative abundance (compared to wildtype) after recovery from the host (R) compared to its abundance in the inoculum (I).

To our surprise, all six tested regulators exhibited significant effects on *C. albicans* colonization fitness. The results for two independent isolates of each mutant are shown in **Figure 1C** and Figure S2. *wor2, wor3, wor4,* and, to a lesser degree, *ahr1* exhibit hypercompetitive phenotypes (**Figure 1C** and Figure S2A, B, C, and D), suggesting that Wor2, Wor3, Wor4, and Ahr1 function as inhibitors of commensalism. Conversely, *czf1* and *ssn6* exhibit diminished fitness compared to wild type, suggesting that Czf1 and Ssn6 promote commensal fitness (**Figure 1C** and Figure S2D). We note that the requirement for Ssn6 in the gut colonization model may not be specific to this niche, given our observation that *ssn6* mutants also exhibit growth defects under multiple *in vitro* conditions (Figure S2F).

The finding that all eight (including Wor1 and Efg1) of eight tested mating factors exert substantial effects on gut colonization suggested that sex and gut commensalism may be controlled by a common regulatory circuit. In support of this hypothesis, Wor1, Efg1, Czf1, and Ahr1 play consistent roles, with each factor serving to either inhibit or promote both processes (**Figure 1D**). In contrast, Wor2, Wor3, Wor4, and Ssn6 display divergent effects on mating and commensalism (**Figure 1D**).

### Genetic epistasis reveals functional relationships among transcriptional regulators in the gut

To clarify the relationships among these sexual regulators in the regulation of commensalism, we turned to genetic epistasis analysis. This classical technique is used to infer the nature of regulatory pathways based on the phenotypes of strains that carry two mutations whose individual phenotypes are distinct. For example, imagine that Factor A promotes a process and Factor B inhibits it. If A and B participate in a linear regulatory pathway (**Figure 2A, linear**), then the loss-of-function phenotype of the more downstream regulator will predominate (be epistatic) in a strain that lacks both factors. Conversely, if A and B are related by parallel or branched regulatory pathways (**Figure 2A, parallel** and **branched**), the phenotype of a double mutant will typically differ from (often assuming an intermediate phenotype between) those of either single mutant.

**Figure 2.**
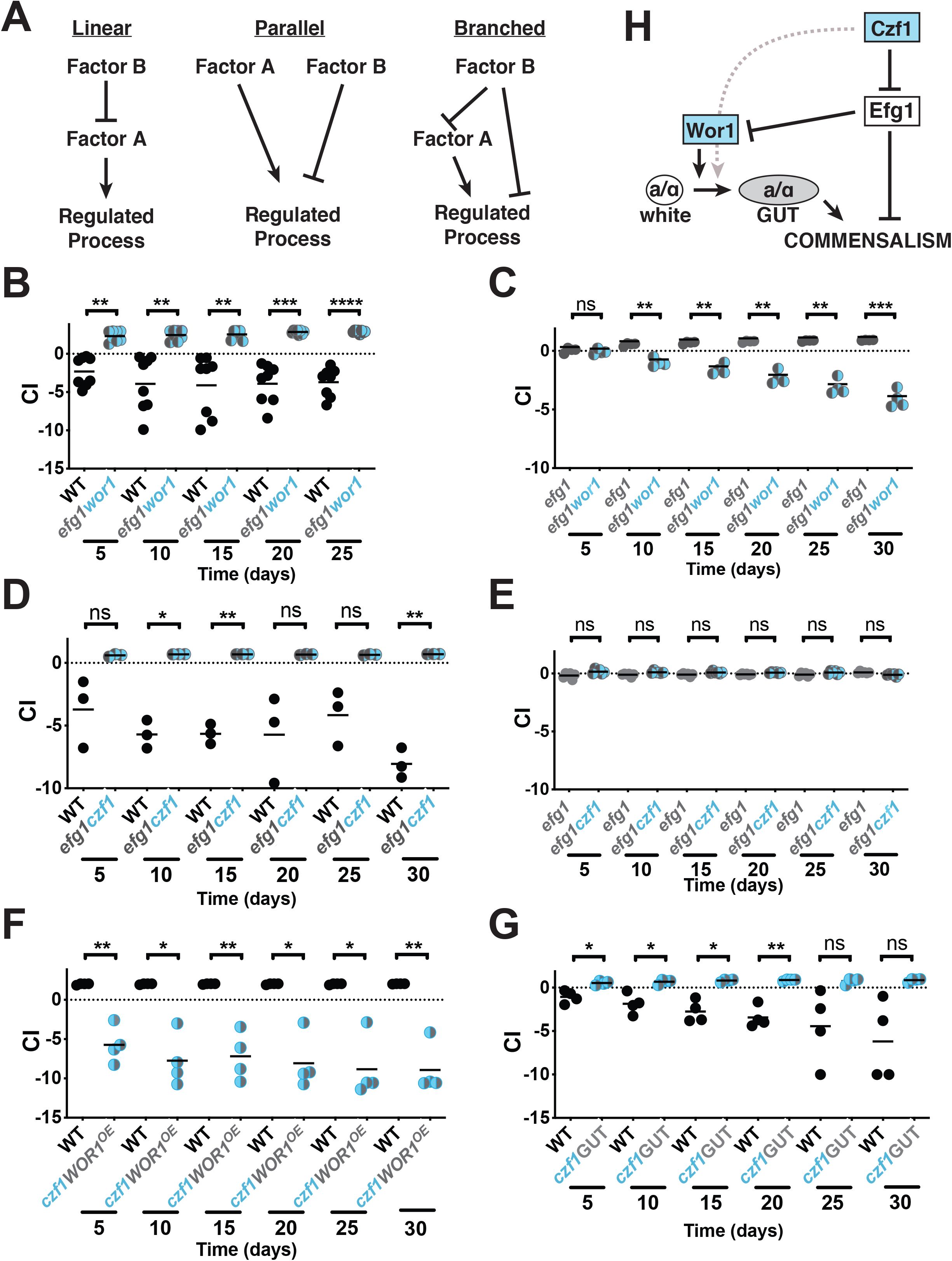
Epistasis analysis reveals a branched regulatory pathway. A. Examples of linear, parallel, and branched regulatory pathways. In Factor A and Factor B have opposite functions in a linear regulatory pathway, then the double mutant *ab* will exhibit the same phenotype as a null mutant affecting the more downstream factor (the diminished output phenotype of mutant *a,* in this example). Alternatively, if Factor A and Factor B participate in parallel or branched regulatory pathways, then the double mutant phenotype *ab* is likely to be intermediate between those of *a* and *b.* B-G. Competitive fitness of selected double mutants in the gut colonization model. B. WT (ySN425) v. *efg1wor1* (ySN1126). C. *efg1* (ySN1338) v. *efg1wor1* (ySN1126). D. WT (ySN425) v. *efg1czf1* (ySN1373). E. *efg1* (ySN1338) v. *efg1czf1* (ySN1373). F. WT (ySN425) v. *czf1WOR1^OE^* (ySN1147). Statistical comparisons were performed using the paired student’s t-test, *p<0.05, ** p<0.01, *** p<0.001, **** p<0.0001. G. WT (ySN425) v. *czf1GUT* (ySN1486). H. Model of commensal regulation inferred from epistasis experiments.

We began with Efg1 (a commensalism inhibitor) and Wor1 (a commensalism promoter). To create an *efg1wor1* double mutant, both copies of *WOR1* were disrupted in an existing *efg1* mutant (note that the *C. albicans* genome is diploid). As shown in **Figure 2B**, an *efg1wor1* double mutant outcompetes wild-type *C. albicans* in the gut colonization model. Nevertheless, it appeared to us that the enhanced fitness of *efg1wor1* is less pronounced than our previous observations with *efg1* single mutants (5,6). To clarify the relative fitness of *efg1* and *efg1wor1*, we performed a direct competition between the two strains in the same animals. As shown in **Figure 2C**, *efg1* consistently outcompetes *efg1wor1*. This result suggests that the magnitude of commensal inhibition by Efg1 exceeds the positive contribution provided by Wor1. Moreover, the finding that an *efg1wor1* double mutant manifests an intermediate phenotype between those of *efg1* and *wor1* single mutants suggests that Wor1 and Efg1 control commensalism by a nonlinear pathway.

We next turned to the relationship between Efg1 and Czf1 (a commensalism promoter). As shown in **Figure 2D**, *efg1czf1* is hypercompetitive in the gut colonization model, similar to the phenotype of *efg1.* Unlike the previous result with the *efg1wor1* mutant, however, *efg1czf1* and *efg1* exhibit identical commensal fitness, with neither strain outcompeting the other in a co-colonization experiment (**Figure 2E**). *EFG1* is thus epistatic to *CZF1*, which suggests that Efg1 lies downstream of Czf1 in a linear regulatory pathway.

Because Wor1 and Czf1 are both activators of commensalism with similar (loss-of-fitness) null mutant phenotypes, we paired a *WOR1*^OE^ gain-of-fitness allele with *czf1* deletions for epistasis analysis. In competition with wildtype, *czf1WOR1*^OE^ exhibits reduced commensal fitness (**Figure 2F**), similar to the phenotype of *czf1* (**Figure 1C**). These results would seem to place *CZF1* downstream of *WOR1* in a linear regulatory pathway. However, as described in the Introduction, Wor1 promotes commensalism by fostering a switch to the GUT cell type, which normally takes between 5 and 10 days. Because the *czf1WOR1*^OE^ strain is outcompeted by wild-type *C. albicans* within the first 5 days of co-colonization, the mutant does not persist for long enough to expose the fitness advantage of the *WOR1*^OE^ allele. To determine whether *czf1WOR1*^OE^ is capable of switching to GUT, we performed monotypic infections of three animals using a pure culture of *czf1WOR1*^OE^ yeasts. Despite stably colonizing the murine GI tract (median ~5×10^7^ CFUs/gram feces), the *czf1WOR1*^OE^ strain yielded no fully GUT colonies over a 30-day time course. This suggests that Czf1 is required for efficient switching to the GUT phenotype. Interestingly, however, when homozygous *czf1* deletions were engineered into a *WOR1*^OE^ strain that had previously been passaged through an animal, the GUT cell phenotype was maintained. Therefore, Czf1 is not required to maintain the GUT phenotype once it has already been established. Moreover, as shown in **Figure 2F**, a GUT phase *czf1WOR1*^OE^ strain outcompetes wild-type *C. albicans* in the murine model, although not to the same extent as *WOR1*^OE^ (5). Together, these results indicate that Czf1 promotes the initiation of the white-to-GUT switch but not maintenance of the GUT phenotype, and that Czf1 and Wor1 are related by a nonlinear regulatory pathway.

The epistasis experiments support a model presented in **Figure 2H**. Czf1 promotes commensal fitness primarily by inhibiting Efg1. The pro-commensalism activity of Czf1 and anti-commensalism activity of Efg1 are inferred from the phenotypes of the respective null mutants, and the ordering of Efg1 downstream of Czf1 is inferred from the enhanced fitness phenotype of *efg1czf1,* which is as fit as the *efg1* single mutant. Efg1, in turn, is posited to inhibit commensalism by a branched pathway, including one arm that involves inhibition of Wor1. The existence of a Wor1-independent mechanism of inhibition is inferred from the ability of an *efg1* mutant to outcompete an *efg1wor1* double mutant in the murine commensalism model. Wor2 and Wor3 also inhibit commensalism, but epistasis experiments with these factors (data not shown) produced complex results that preclude their inclusion in the model.

We previously showed that Wor1 fosters commensalism by promoting a white-to-GUT transition to a commensal-specific cell type (5). Czf1 is required for switching to occur, either by directly promoting the switch (as depicted in **Figure 2H**) or by inhibiting Efg1. This is supported by the failure of *czf1WOR1*^OE^ white cells to switch to GUT during a monotypic colonization experiment. However, Czf1 is not required to maintain the GUT phenotype, once established, given our ability to generate *czf1WOR1*^OE^ GUT cells with moderately enhanced fitness in the commensal model. Continued expression of Efg1 in the absence of Czf1 likely accounts for the failure of this strain to exhibit the more pronounced gain-of-fitness phenotype of a *WOR1*^OE^ GUT strain.

We note that the core genetic logic of the commensalism pathway recapitulates that of the sexual circuit, with Czf1 inhibiting Efg1, Efg1 inhibiting Wor1, and Wor1 promoting a cell type switch that promotes the regulated process. Apparent differences include a proposed role for Czf1 in promoting (and/or for Efg1 in inhibiting) the white-to-GUT switch, as well as a Wor1-independent branch of Efg1-mediated repression of commensalism.

### Calling Card-seq records transcription factor binding events in *Candida albicans*

To further explore the activity of these regulators, we sought a method to determine their DNA binding activities in commensally propagated cells. Chromatin immunoprecipitation (ChIP) is a widely used technique for identifying the direct DNA binding targets of transcription factors. However, because this biochemical technique utilizes relatively large numbers of cells, the strains are typically propagated in the laboratory under conditions that foster activity of the relevant transcription factors. Because there are no known *in vitro* conditions that adequately mimic the lumen of the mammalian gastrointestinal tract, we developed an alternative strategy that would permit the localization of transcription factor binding sites in cells propagated within the host.

Originally developed for use in *S. cerevisiae* and later in mammalian cells, the Calling Card-Seq technique utilizes a copy of PiggyBac transposase (PBase) that is fused to a transcription factor (TF) of interest to deposit copies of PiggyBac transposon (PB) into TTAA sequences that abut the binding sites of the TF (20,21). Transposition is mediated within living cells, and binding sites are determined by DNA sequencing of transposon insertion sites across a cell population. Multiple modifications were required to adapt this technique for use in *C. albicans* (**Figure 3A**, Figure S3, and Methods). First, until very recently (22), there have been no stable autonomously replicating plasmids available for use in *C. albicans.* We therefore integrated one or two copies of PB into a neutral locus in the *C. albicans* genome. Second, because *C. albicans* uses a non-universal genetic code (CTG is used to encode leucine rather than serine; (23)), we synthesized a codon-optimized version of PBase. Finally, to facilitate monitoring and selection of transposon excision and reinsertion events, we utilized Arg^−^His^−^Leu^−^ reference strain SN152 as the parent strain for these studies. To permit monitoring of transposon excision, 5’ and 3’ halves of *ARG4* were placed on either side of PB at the chromosomal donor site, such that precise excision of the transposon results in prototrophy for arginine. To monitor and select for transposon reinsertion, an intact copy of *HIS1* was embedded within PB, such that cells that have undergone transposon excision and reinsertion are prototrophic for both arginine and histidine. For experiments in animals, we used a Calling Card strain that contains two transposons, one marked with *HIS1* and the second marked with *LEU2.*

**Figure 3.**
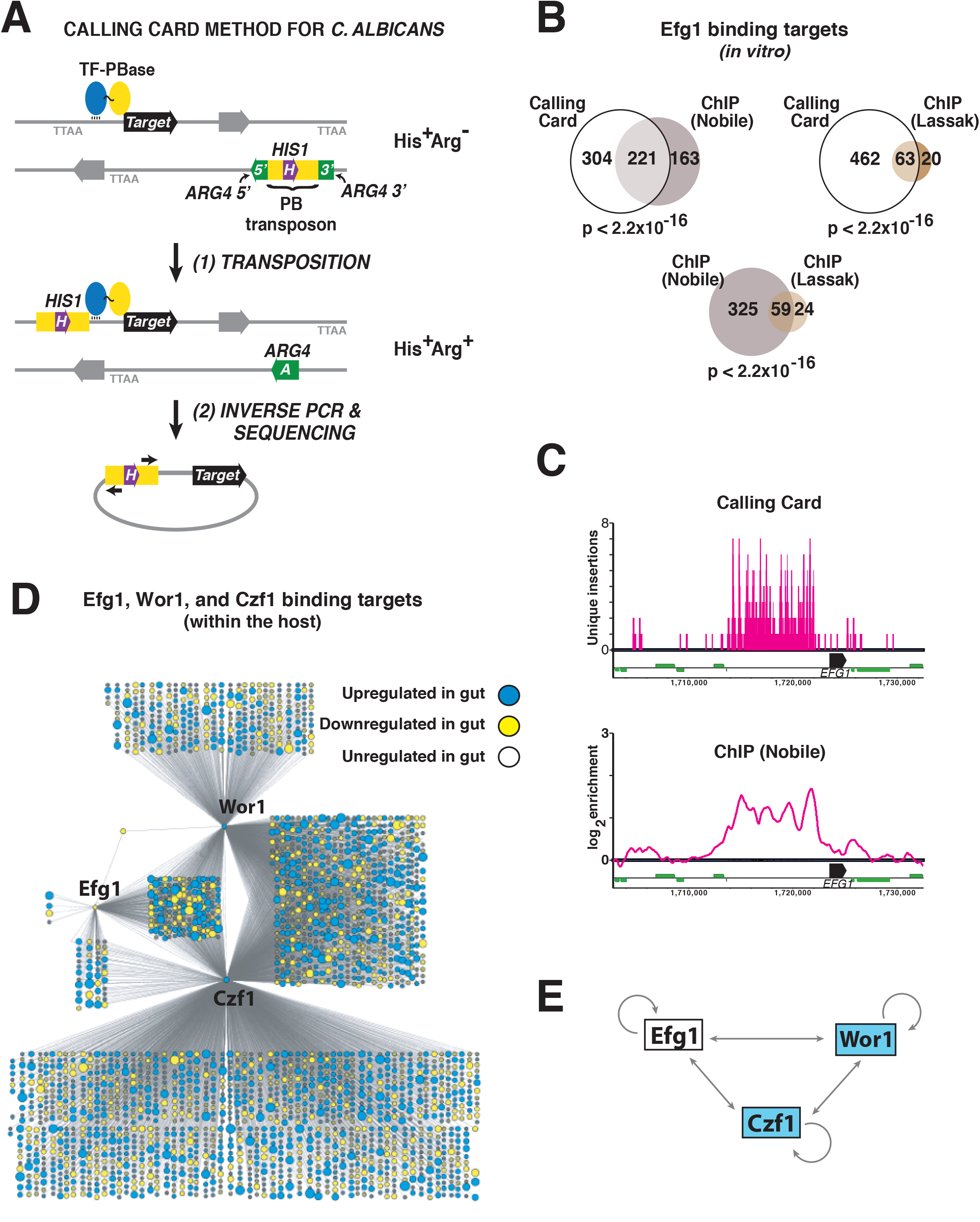
Determination of transcription factor binding sites *in vitro* and in the host using Calling Card-Seq. A. Cartoon of the Calling card-seq method. Test strains contain one or two copies of PiggyBac transposon (PB) and express a transcription factor (TF)-PiggyBac transposase (PBase) fusion protein. Propagation of these strains in the laboratory or in a mammalian host allows for PBase-mediated excision of PB from a neutral donor site and reinsertion it into a TTAA sequence abutting the binding site of the TF. Because PB at the original donor site interrupts a copy of *ARG4,* transposon excision confers an Arg+ phenotype. Because PB contains the *HIS1* gene, cells that have undergone transposon reinsertion exhibit an Arg+His+ phenotype (or Arg+His+Leu+ in cells that contain two copies of PB). B. Similar Efg1 binding sites are determined by Calling Card-Seq and ChIP. Venn diagrams depict the overlap between the 525 high confidence binding Efg1 binding sites identified using Calling Card-Seq and the 83 and 384 binding sites previously identified by Lassak et al. (25) and Nobile et al. (26) using ChIP. *In vitro* cell culture conditions for Calling Card-Seq were Lee’s 2% GlcNAc/agar pH 6.8, 25°C; Lassak YPD, 30°C (25); and Nobile Spider, 37°C (26). Significance of target overlap was determined by Fisher’s exact test. C. Comparison of results obtained using Calling Card-Seq vs. ChIP r for Efg1 binding to the *EFG1* locus. Calling card-Seq insertion events are depicted in the upper panel, ChIP binding peaks reported by Nobile et al. (26) are depicted on the lower panel. D. Regulatory networks of Efg1, Wor1, and Czf1 during commensal growth in the host. Efg1-PBase, Wor1-PBase, and Czf1-PBase strains were propagated in the murine gut commensalism model, and high confidence direct binding targets were determined using Calling Card-Seq. Each node corresponds to a direct binding target of Efg1, Wor1, and/or Czf1, with node size reflecting the fold-change in expression of the corresponding ORF when WT *C. albicans* cells is propagated in the gut colonization model (compared to *in vitro* growth in YPD, 30°C (6)). Node color indicates upregulation (yellow), downregulation (blue), or no change (empty circle of fixed size) in the gut vs. *in vitro*. E. Schematic of cross regulation among Efg1, Wor1, and Czf1 within the host digestive tract. Results obtained with Calling Card-Seq suggest that, unlike under *in vitro* conditions, **a**/α cells express all three regulators within the mammalian gut.

To test the modified Calling Card system, we deployed it to map the binding sites of an Efg1-PBase fusion protein in *C. albicans* yeasts that were propagated *in vitro*. A strain encoding untethered PBase under regulation of the *EFG1* promoter was used as a control (*n.b.* PBase contains its own nuclear localization signal (24)). Transposition in both strains was allowed to occur for seven days, during propagation on solid medium (Lees 2% GlcNAc/agar pH 6.8) maintained at 25°C. Colonies containing transposon reintegrants were selected by replica plating to medium lacking arginine and histidine. Genomic DNA was extracted from Arg^+^His^+^ colonies, and genomic DNA extraction, inverse PCR, and sequencing library preparation was performed as described in the Methods. Transposon insertion sites were mapped to the genome, and background signal associated with the untethered control was subtracted from the Efg1-PBase signal. Finally, high confidence hits for Efg1-PBase were defined as promoters associated with >5 independent transposon insertions (i.e. at different TTAA sequences) within a 1000 bp window and p<0.05 using a Poisson or hypergeometric test.

The modified Calling Card-Seq technique identified 525 high confidence DNA-binding targets, as detailed in Table S1. Two groups have previously used ChIP to define Efg1 binding targets in **a**/α strains. Lassak et al. reported 83 Efg1 binding sites in *C. albicans* yeasts propagated in YEPD medium at 30°C (25), and Nobile et al. 384 Efg1 binding sites in *C. albicans* biofilms propagated in Spider medium at 37°C (26). Notably, the set of targets identified using Calling Card-Seq exhibits significant overlap with both sets of ChIP-defined targets (63 in common with the Lassak dataset, p<2.2×10^−16^ and 221 in common with the Nobile dataset, p<2.2×10^−16^). Moreover, the significance of overlap between Calling Card-defined targets and either set of ChIP-defined targets is approximately the same as that between the two sets of ChIP-defined targets (59 common targets, p<2.22×10^−16^; **Figure 3B**).

### Recording of transcription factor binding events during mammalian gut commensalism

To determine the activities of Efg1, Wor1, and Czf1 during commensalism, we utilized strains containing Efg1-PBase, Wor1-PBase, or Czf1-PBase. Each strain was gavaged into an independent group of BALB/c mice (n=4-5 animals per strain), and colonization was allowed to progress for 15 (Efg1-PBase and Czf1-PBase) or 20 (Wor1-PBase) days. Animals were euthanized periodically over the experimental time course, and yeasts recovered from ceca (the gut compartment containing the highest burden of *C. albicans*) were plated onto medium lacking arginine, histidine, and leucine to select for transposon excision and reintegration events. An untethered PBase-overexpression strain (PBase^OE^) was propagated *in vitro* to serve as a negative control. Genomic DNA extraction, inverse PCR, and preparation of sequencing libraries was performed as described in the Methods.

Following subtraction of background signal (signal from untethered PBase) and after thresholding for high confidence hits, Calling Card-seq revealed 2561 *C. albicans* genes that are targeted by at least one of the transcription factors in commensally propagated cells (**Figure 3D** and Table S2). Of note, Wor1, Czf1 and Efg1 each bind to their own promoters as well as to the promoters of the other two transcription factors (**Figure 3D** and **3E**). Similar results for Efg1 have been reported by Nobile et al., based on ChIP experiments using cells propagated under *in vitro* conditions (26); however, binding targets of Wor1 and Czf1 have not previously been investigated in **a**/α cells. Indeed, it would be difficult to analyze Wor1 using ChIP because of its very low abundance in this cell type under *in vitro* conditions. The *in vivo* Calling Card analysis revealed 177 genes that are targeted by all three DNA-binding proteins (**Figure 3D** and Table S2), including ones encoding 24 other transcription factors. Importantly, eight of these transcription factors have demonstrated roles in the regulation of commensalism: Wor2, Wor3, Wor4, and Ahr1 (shown in **Figure 1**), as well as Tec1, Brg1, and Ume6 (6) function as inhibitors of gut colonization, whereas Crz2 has been reported to enhance the establishment of colonization (27). These experiments, which record transcription factor activity during host colonization, reveal the DNA binding activities of Wor1, Czf1, and Efg1 in the commensal niche, which includes a large set of common regulatory targets.

### Comparative transcriptomics of circuit-altered strains uncovers commensalism effectors

Our previous unbiased screens for commensalism factors have exclusively identified genes encoding transcriptional regulators (5,6), likely reflecting the outsized effects of proteins that regulate the expression of hundreds or even thousands of genes. We hypothesized that regulatory mutants with contrasting commensal phenotypes might serve as tools for the discovery of commensalism effectors. Specifically, we hypothesized that comparison of the RNA expression patterns of strains with enhanced vs. diminished commensal fitness would reveal genes mediating direct interactions with the host and co-colonizing microorganisms.

To test this hypothesis, we used RNA-Seq to determine the transcriptomes of *C. albicans* mutants with hyperfit vs. hypofit phenotypes in the gut colonization model. Groups of BALB/c mice (n=5-6 per strain) were colonized with *wor1, czf1*, *efg1*, *WOR1*^OE^ GUT cells, *efg1wor1*, or wild-type *C. albicans*. After 10 days of propagation in the host, strains were recovered from mouse large intestines, and RNAs were extracted for RNA-Seq. For comparison, the strains were also profiled under *in vitro* conditions (YEPD, 30°C).

Shown in **Figure 4A** is a volcano plot of results for strains with enhanced (*efg1*, *WOR1*^OE^ GUT cells, *efg1wor1*) vs. diminished (*wor1, czf1*) commensal fitness. For each gene, relative RNA abundance (log_2_[abundance in hyperfit/abundance in hypofit]) is plotted on the x-axis, and significance (-log_10_[adjusted p-value]) is plotted on the y-axis; all RNA-Seq results are presented in Table S3. In total, 312 genes were significantly (p<0.05) upregulated by 2-fold or more in the hyperfit strains, whereas 590 were downregulated. We hypothesized that genes that inhibit commensal fitness would be downregulated in hyperfit strains. In support of this idea, GO-term analysis of the downregulated gene set revealed significant enrichment for hypha-associated genes (hyphal cell wall, p=6.48×10^−7^). In previous work, we have shown that hypha-specific cell wall and secreted factors diminish the fitness of *C. albicans* in the gut colonization model, presumably because of host restriction of this invasive cell type (6).

**Figure 4.**
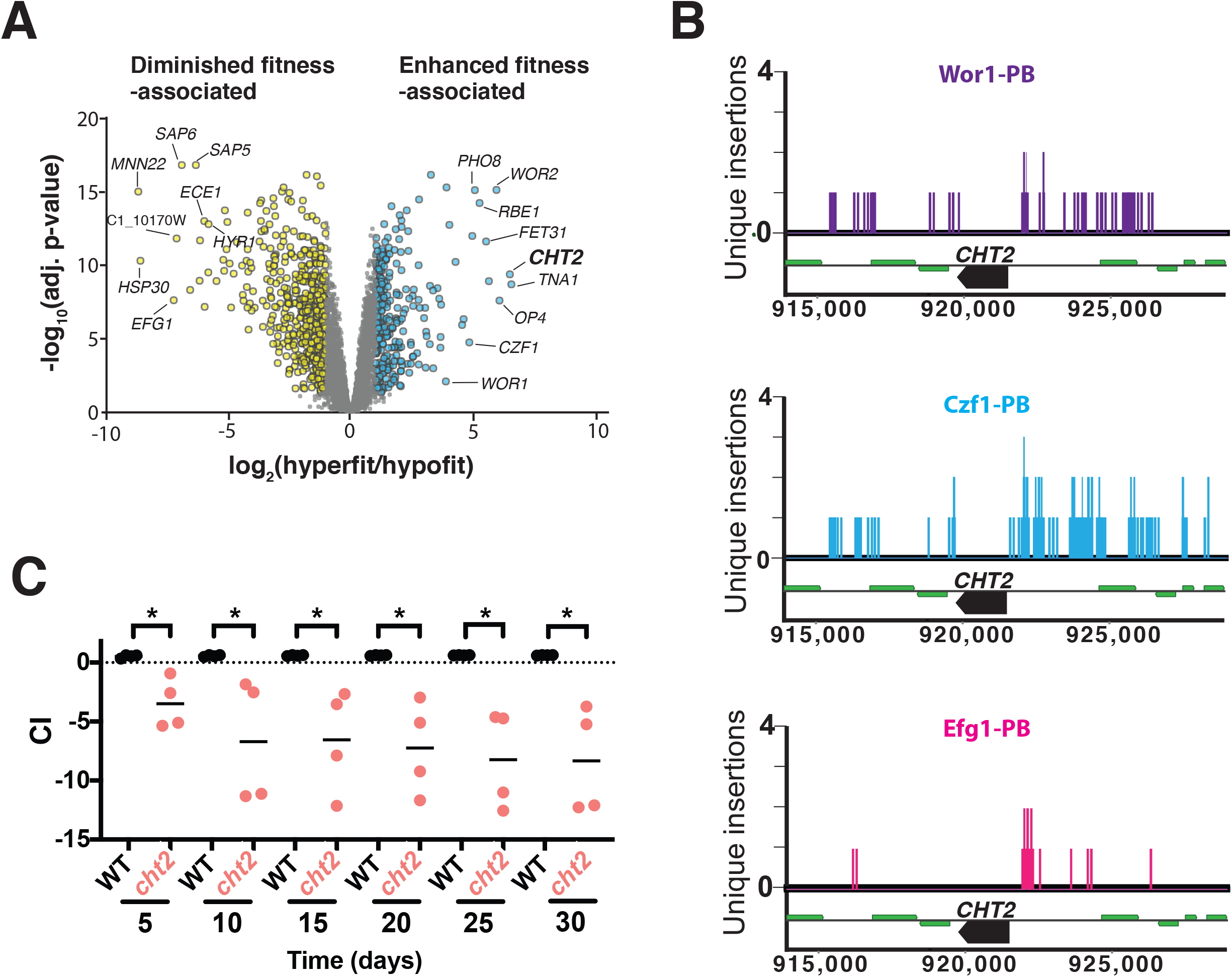
The transcriptomes of transcription factor mutants displaying enhanced vs. diminished fitness reveal effectors of commensalism. **A.** Volcano plot of the log_2_ transformed ratio of gene expression among mutants with enhanced commensal fitness (*efg1*, *WOR1*^OE^ GUT cells, *efg1wor1*) compared to those with commensal defects (*wor1, czf1*; x-axis) versus significance (y-axis). **B.** Calling Card reveals direct binding of Wor1, Czf1, and Efg1 to the *CHT2* promoter in commensally growing cells. **D**. *CHT2* is required for normal fitness in the murine gut colonization model. Wild type (ySN250) and *cht2* (ySN2104) were competed in four animals. *p<0.05.

We hypothesized that genes upregulated in hyperfit strains would include ones for commensalism effectors. GO-term analysis of this gene set was less informative, showing significant enrichment of genes associated with processes such as translation (ribosome, p=4.51×10^−8^) and sugar transport (monosaccharide transmembrane transporter activity, p=0.00119), among others. To determine whether any of these genes promotes commensal fitness, we focused on *CHT2,* a gene that encodes a GPI-linked fungal chitinase, which is substantially (6.4-fold) and significantly (p<0.001) upregulated in the hyperfit strains. In addition, our Calling Card-Seq data indicate that the *CHT2* promoter is directly bound by Efg1, Wor1, and Czf1 in commensally propagated cells (**Figure 4B**). A *cht2* homozygous gene deletion mutant was constructed and tested in the mouse gut colonization model. As shown in **Figure 4C**, *cht2* is outcompeted by wildtype, suggesting that this factor is required for normal fitness in the gut.

## DISCUSSION

In this work, we establish roles for 8 key regulators of *C. albicans* mating in the regulation of fungal commensalism. Null mutants affecting *WOR2, WOR3, WOR4, AHR1, CZF1,* and *SSN6* exhibit altered fitness in a mouse model of gut colonization, as we have previously demonstrated for mutants affecting *WOR1* and *EFG1* (5). Moreover, genetic epistasis analysis of double mutants assayed in the host indicates that core relationships among Czf1, Efg1, and Wor1 are preserved between sex and host colonization. The repurposing of a complex regulatory pathway for two very different outcomes was unexpected. However, our *in vivo* analysis of transcription factor DNA binding activity using Calling Card-Seq supports activity of all tested regulators in commensally growing yeasts.

The development of a modified Calling Card-seq technique for *C. albicans* enables the study of transcription factor binding activity during infection, rather than under laboratory conditions. Animal models exist for diverse *C. albicans* interactions with the host, including commensal and pathogenic interactions with skin, with the oropharynx, with the vagina, and with the gut, in addition to multiple animal models of systemic infection (28–35). Supporting the validity of the Calling Card-Seq method, our results for Efg1-PBase under *in vitro* conditions demonstrated significant overlap with results previously reported using ChIP (25,26), and results for the *in vivo-*propagated Efg1-PBase, Czf1-PBase, and Wor1-PBase strains identified multiple targets with known roles in commensalism.

Having identified sets of strains with enhanced versus diminished commensal fitness, we utilized comparative transcriptomics to identify potential mediators of fitness. In keeping with our previous observations that certain hypha-associated genes inhibit the commensal fitness of *C. albicans* in the gut (6), genes encoding hypha-associated cell wall proteins were enriched among the set of downregulated genes in hyperfit strains. Meanwhile, we discovered that *CHT2* gene for a GPI-linked fungal chitinase, which is induced in hyperfit strains, promotes commensal fitness in the gut colonization model. As chitin is present in the fungal cell wall, in bacteria, and in the diet but not in mammalian tissues, it remains to be determined how this chitinase functions in commensalism.

Mating and the white-to-opaque epigenetic switch are among the most highly studied processes in the *C. albicans* field. Nevertheless, there is little evidence for sexual exchange among naturally circulating strains (36,37), and mating is inferred to be exceedingly uncommon. In contrast, gut colonization with *C. albicans* is highly prevalent among humans and other mammals, and we speculate that propagation within this commensal niche may provide ongoing selection for circuit components that are shared between mating and commensalism. Further work will be required to determine how the same transcriptional regulators can direct two very different outcomes (mating vs. host colonization).

## ACKNOWLEDGEMENTS

We are grateful to Rob Mitra, Zongtai Qi and Xuhua Chen for generously sharing plasmids and software that they developed for the mammalian piggyBac transposase system. We thank Hiten Madhani, Rob Mitra, and Anita Sil for helpful comments on the manuscript. This work was supported by US National Institutes of Health (NIH) grant R01AI108992, a Burroughs Welcome Award in the Pathogenesis of Infectious Disease, and a Pew Foundation scholarship. JNW was supported by NIH grant T32AI060537 and a Discovery Fellows Grant from UCSF.

## MATERIALS AND METHODS

The yeast strains used for this study are described in Table S4, plasmids in Table S5, and primers in Table S6.

### Studies in animals

All procedures involving animals were approved by the UCSF Institutional Animal Care and Use Committee. The mouse model of *C. albicans* commensalism was performed as previously described (5,38) with minor modifications. Groups of 8–10-week (18-21 gram) female BALB/c mice were housed 2 per cage, unless otherwise specified, and treated with penicillin 1500 un/ml and streptomycin 2 mg/ml in drinking water starting 7 days prior to gavage, and continuously for the remainder of the experiment. Feces were plated on Sabouraud agar, LB agar, and MRS agar to monitor for fungal and bacterial growth.

For commensal competition experiments, groups of 4-8 mice were gavaged with 10^8^ CFUs of a 50:50 mix of two fungal strains in 0.9% saline. *C. albicans* was recovered from the inoculum and host feces at specified intervals, and strain quantification was performed as described previously (39).

For mRNA-seq experiments, mice were colonized with a single strain and individually housed. After 10 days, animals were humanely euthanized, and large intestines were dissected for RNA analysis.

### mRNA-Seq library preparation and analysis

RNA extraction for mRNA-seq was performed as previously described (6), with modifications. Luminal contents of large intestines were resuspended in a mixture of 500 μL Buffer A (200 mM NaCl and 20 mM EDTA), 500 μL acid phenol-chloroform, 168 μl 25% SDS, and ~400 μL 0.5 mm glass beads. Samples were vortexed for 5 cycles of 2 minutes, interrupted with 1 minute rests on ice. Samples were centrifuged at max speed in a microcentrifuge for 10 minutes, and supernatants were extracted four times with equal volumes of acid phenol-chloroform (Ambion), followed by four rounds of extraction with equal volumes of phenol-chloroform-isoamyl alcohol (Ambion). Following precipitation and resuspension of pellets in RNase-free water, RNA was further purified using the MEGAclear transcription clean-up kit (Ambion) for *wor1* (ySN1351), or Sera-Mag beads in a homemade PEG solution (https://openwetware.org/wiki/SPRI_bead_mix) following the RNAclean XP protocol for ySN425, *efg1* (ySN1011), *czf1* (ySN1145), *WOR1^OE^* GUT (ySN1045), and *efg1wor1* (ySN1126). DNase I (NEB) treatment was performed on at least 5 µg RNA for 10 minutes at 37°C. Following DNase treatment, a final acid phenol-chloroform extraction was performed. RNA was then precipitated and resuspended in RNase-free water.

NEBNext Ultra Directional RNA Library Prep Kit for Illumina was used in combination with NEBNext Poly(A) mRNA Magnetic Isolation Module and NEBNext Multiplex Oligos for Illumina to generate mRNA-seq libraries. The protocol provided by NEB was followed using 1 µg total RNA and 14 cycles of PCR, with the exception that Ampure XP bead mix was replaced with Sera-Mag Speed Beads (Thermo-Fisher) in a homemade PEG solution (40). Library fragment size was determined using High Sensitivity DNA chips on a 2100 Bioanalyzer (Agilent). Library quantification was performed by qPCR with a library quantification kit from KAPA Biosystems (KK4824). Sequencing was performed on a HiSeq4000 device (Illumina).

Sequencing reads were mapped to the current haploid *C. albicans* transcriptome (Assembly 22 default coding, downloaded from candidagenome.org on November 19, 2018) and transcript abundances were determined using kallisto (41). Statistical comparisons of transcript abundances between different samples were performed on estimated counts generated by kallisto using *limma* as previously described (42). The analysis of hyperfit vs. hypofit transcriptomes was performed by binning RNA expression results of strains with enhanced vs. diminished commensal fitness relative to wild-type *C. albicans,* and using limma to compare the two groups.

GO term analysis was performed using the Candida Genome Database GO term finder. Differentially expressed genes were considered to be those with a fold change greater than 2 and an adjusted p-value less than 0.05. Upregulated or downregulated genes were analyzed for process, function and component.

### Generation of *C. albicans* strains for Calling Card-Seq

Because, stable, autonomously replicating plasmids have not been available for use in *C. albicans* until quite recently (22), we devised a strategy to integrate recombinant Piggyback transposons, genes encoding transcription factor-PBase fusion proteins, and codon-optimized *PBASE* into the genome. First, we commissioned synthesis of a PBase gene (*PBASE*) that is codon-optimized for *C. albicans* (GenScript, Piscataway, NJ). Next, we engineered a series of plasmids to target codon-optimized *PBASE* to the 3’ end of a transcription factor ORF of interest; integration results in expression of a TF-PBase fusion protein via the natural promoter for the TF. Plasmid pSN427 (which is used to create TF-specific integrating constructs) contains the following insert: PmeI restriction site—sequence encoding an 18 amino acid linker— *PBASE*—*FLP*—*SAT1*—PmeI restriction site. pSN427 was generated by homologous recombination in *S. cerevisiae* of the following three PCR products: 1) Codon-optimized *PBASE,* amplified with primers that include linker homology and a PmeI site at the 5’ end and homology to *FLP-SAT1* cassette at the 3’ end. 2) Linearized pRS316 vector, amplified with primers containing linker homology on one end and *FLP-SAT1* cassette homology on the other. 3) *FLP-SAT1,* amplified from pSFS2A using primers with homology to *PBASE* at the 5’ end and pRS316 at the 3’ end.

To create custom integration constructs that target *PBASE* to the *WOR1* (pSN459)*, EFG1* (pSN467), and *CZF1* (pSN465) ORFs, homologous recombination in *S. cerevisiae* was used to fuse the following three inserts: 1) linker—*PBASE*—*FLP*—*SAT1* fragment from PmeI-digested pSN427; 2) PmeI restriction site—terminal ~300 bp of transcription factor ORF sequence; 3) 300 bp of 3’ UTR for the transcription factor—PmeI site. To create a PBase^OE^ control strain, an analogous method was used to create pSN422, which targets *PBASE* to replace one copy of the *TDH3* ORF. pSN422 contains the following insert: PmeI—~300 bp of *TDH3* 5’ UTR—linker—*PBASE*—*FLP*—*SAT1*—~300 bp of *TDH3 3’* UTR-PmeI.

Inserts from the PBase^OE^ (pSN422) and transcription factor-PBase plasmids (pSN459, pSN467, pSN465) were liberated with PmeI and used to transform *C. albicans* strains containing either one (ySN1873) or two (ySN1877) copies of PB transposon (at the neutral *leu2*∆ locus). Following 1 day of propagation on YPD/agar containing 200 ug/ml nourseothricin (YPD+Nat) at 30°C, Nat-resistant colonies were streaked to fresh YPD+Nat medium and incubated overnight at 30°C. Finally, single colonies were patched to YPD+Nat, SD-histidine, SD-histidine-arginine, and SD-histidine-leucine-arginine plates. 5’ and 3’ junctions of integrants were verified by colony PCR using primers described in Table S6. Glycerol stocks were prepared from patches exhibiting His^+^ (1 transposon) or His^+^Leu^+^ (2 transposons) and Arg^−^ (transposon still present at the donor site) phenotypes. In addition, we screened for Nat^S^ versions of the strains following overnight propagation in YPD+maltose liquid culture at 30°C, which allows for spontaneous excision of the *FLP-SAT1* cassette.

To validate the modified method under *in vitro* conditions, Efg1-PBase and *EFG1* promoter-PBase strains (containing one copy of PB) were propagated for 2 days on YEPD, followed by replating of ~50,000 onto Lee’s GlcNAc pH 6.8 agar (hypha-inducing conditions where *EFG1* is well-expressed) in 100 mm petri plates. Strains were incubated at 25°C for 7 days to allow for transposition to occur. Cells were then replica-plated to SD-histidine-arginine and incubated for 2 days at 30°C. His+Arg+ cells were collected by washing with water, followed by extraction of genomic DNA from cell pellets.

### Calling Card-Seq mouse experiments

*WOR1-PBASE* (ySN1933), *EFG1-PBASE* (ySN1943), and *CZF1-PBASE* (ySN1941) strains containing two copies of PB were streaked from frozen glycerol stocks onto YPD plates and incubated for 16 hours at 30°C. Cells were scraped from the plates, diluted to 5×10^5^ CFU/ml, and 100 μl each cell suspension was spread onto several SD-histidine-leucine plates, with incubation for 48 hours at 30°C. A disposable yellow loop was used to scrape colonies into 0.9% saline. 10^8^ CFUs was gavaged into each antibiotic-treated mouse. One gavage volume was plated to SD-histidine-leucine-arginine plates to monitor for transposition events that had already occurred during preparation of the inocula (these proved to be minimal; Witchley and Noble, data not shown). To estimate the background rate of nonspecific transposition events, the PBase^OE^ (ySN1904) was grown in liquid YPD or YPD+100 μM BPS (iron chelator related to TFs not reported here) overnight and then plated to selection media.

After 5, 10, 15, and/or 20 days, animals were euthanized and dissected using sterile technique to recover the ceca. 1-2 animals per group were euthanized after 5, 10, or 15 days for Efg1-PBase, 10 or 15 days for Czf1-PBase and 10, 15 and 20 days for Wor1-PBase. Each organ was homogenized in 3 ml PBS pH 7.4. 100 μl was diluted 1:10 in PBS to determine CFUs/gram cecum, and the remainder was plated 500 μl per 100 mm SD-histidine-leucine-arginine plate. Cells were incubated for 2 days at 30°C, and colonies were recovered for DNA extraction and sequencing library preparation. All reads from the same strain recovered from animals were pooled prior to data analysis.

### Calling Card sequencing library preparation and analysis

Sequencing libraries were prepared as previously described (20,21) with modifications. Genomic DNA was digested MspI, RsaI and Taq1α, in independent reactions. Next, the DNA was precipitated with isopropanol and 5 M ammonium acetate, washed with 70% ethanol and resuspended in 30 μl nuclease-free water. To circularize the pooled restriction fragments, the DNA was resuspended to 2-3 ng/μl in a 400 μl ligation reaction with T4 ligase. The ligation reaction was incubated at 15°C for greater than 16 hours then precipitated with ethanol and 3 M sodium acetate, washed with 70% ethanol and resuspended in 30 μl nuclease-free water. 15 μl of ligated DNA was used in a PCR reaction with Phusion polymerase and custom read one primer SNO2980 and one of 96 barcoded paired end primer 2. The PCR program used was: Step 1: 94°C 2 min, Step 2: 94°C 30 seconds, Step 3: 94°C 30 seconds, go to Step 2 for 30 cycles, hold at 10°C. PCR cleanup was performed using 1.0x homemade SPRI bead mix (40). PCR product quantification was performed using Quant-iT PicoGreen dsDNA assay kit (P11496). After quantification, all reactions were pooled in equal quantities. Size selection was performed on the Center for Advanced Technology (UCSF) Sage Science BluePippin with 2% agarose dye-free cassettes with marker V1 using 250-500 base pair elution criteria. A subsequent buffer exchange to TE was performed with SPRI bead mix (40). Libraries were visualized using High Sensitivity DNA chips on an Agilent Bioanalyzer. Sequencing was performed on a HiSeq4000 using custom read 1 sequencing primer SNO3189.

Sequenced reads were aligned to *C. albicans* diploid genome Assembly 22 using bowtie (43). Custom Calling Card-seq analysis code provided by the Mitra lab was adapted for use in *C. albicans* as follows: To control for possible jackpots in PCR amplification, all reads mapping to a specific genomic integration position were collapsed to a single instance of integration at that position. To limit random sequencing noise, a minimum of 50 reads per integration site was required to include the site. Transposon insertions were grouped into clusters within 1000 base pair windows. Insertion clusters for a given transcription factor-PBase fusion protein were compared with background signal (i.e. clusters associated with untethered PBase^OE^), and significant differences at each position were determined using Poisson and Hypergeometric distributions. Clusters were annotated according to the names of the two closest genes using bedtools closest-feature. For interpretation, the data were collapsed by assigning clusters to the closest promoters and subtracting all untethered PBase transposition events from the TF-PBase insertions. If insertions occurred between two genes transcribed in opposite directions away from each other, both genes were assigned to these insertions. High confidence transcription factor binding targets were defined as genes associated with at least 5 independent insertions above background and a p-value <0.001 (Poisson or Hypergeometric distribution) for at least one of the clusters upstream of the target gene. These data are presented in **Figure 3B** and **D,** Table S1, and Table S2.

For visualization in MochiView (44), duplicate reads were first removed using a two-step process: duplicates in each BAM (binary sequence alignment map) file were flagged using Picard MarkDuplicates (Broad Institute, MIT); then, if the number of duplicate reads was greater than 50, a single read was kept in the BAM file as the “Unique Insertion”, else the reads were discarded. The genomeCoverageBed function of bedtools (45) was used to collect sequencing depth information in a bedGraph file format. The bedGraph files were then converted to fit the MochiView file format (44). Formatted files were imported into MochiView as a Location/Data set. A bar graph style was used for visualization due to the nature of the data and the MochiView file format. These data are presented in **Figure 3C and Figure 4B.**

**Figure S1.** *C. albicans* **sexual regulators bind to each other’s promoters in white and opaque a cells (Related to Figure 1).** Cartoons of DNA binding activity in white **a** cells (left) and opaque **a** cells (right) are based on the results of ChIP experiments reported by other groups (13,14,16). Each dot indicates a direct binding interaction, and only the boxed regulators are expressed in each cell type. Cells used for ChIP experiments were propagated *in vitro.*

**Figure S2. Validation of commensal phenotypes with independent isolates of***wor2, wor3, wor4, czf1,* **and***ahr1* **(Related for Figure 1).** A-E. Independently generated isolates of each mutant were competed against WT in the murine gut colonization model. A. WT (ySN250) v. *wor2* (ySN1143). B. WT (ySN250) v. *wor3* (ySN1433). C. WT (ySN250) v. *wor4* (ySN1494). D. WT (ySN250) v. *czf1* (ySN1383). E. WT (ySN250) v. *ahr1* (ySN1188). Paired student’s t-test *p<0.05, **p<0.01, ***p<0.001, ****p<0.0001. F. *ssn6* exhibits colony growth and filamentation defects on multiple types of laboratory media. Single colonies of WT and *ssn6* are depicted on YEPD, Lee’s glucose, and Spider plates incubated at 30°C and 37°C.

**Figure S3. Calling card-seq constructs and validation of activity (related to Figures 3 and 4).** A. Schematic of PB transposons at the donor site. *C.d.HIS1-* and, in strains containing two transposons, *C.m.LEU2-*marked PB are flanked by a split *ARG4* gene. The split *ARG4-* transposon constructs are integrated into the chromosomal *leu2*∆ locus (both endogenous allele of the *LEU2* ORF are deleted in the SN152 parent strain). B. Use of auxotrophic markers to monitor transposon excision and reintegration events. Prior to transposase activity, strains containing one copy of PB have a His+Arg-phenotype. Strains that have undergone transposon excision and loss of PB have a His-Arg+ phenotype. Strains that have undergone transposon excision and reintegration into the genome have a His+Arg+ phenotype. C. Schematic of method to create Pbase^OE^ strains The insert from PmeI-digested pSN422 undergoes recombination at the *TDH3* locus, resulting in replacement of the *TDH3* ORF. As a result, untethered PBase is expressed at high levels via the strong, constitutively active *TDH3* promoter. D Growth of strains containing only the PB-*HIS1* transposon (upper half) vs. PB-*HIS1* plus untethered PBase (lower half) on various media. Both strains grow well on YEPD/agar (left, nonselective) and on SD-his/agar (middle, selective for presence of the transposon), but only the strain expressing PBase grows on SD-arg (right; selective for the presence of the transposon and for excision of the transposon from the donor site).

