## Supplementary figures and images for "Recording of DNA binding events during gut commensalism reveals the action of a repurposed *Candida albicans* regulatory network"

### Figure S1

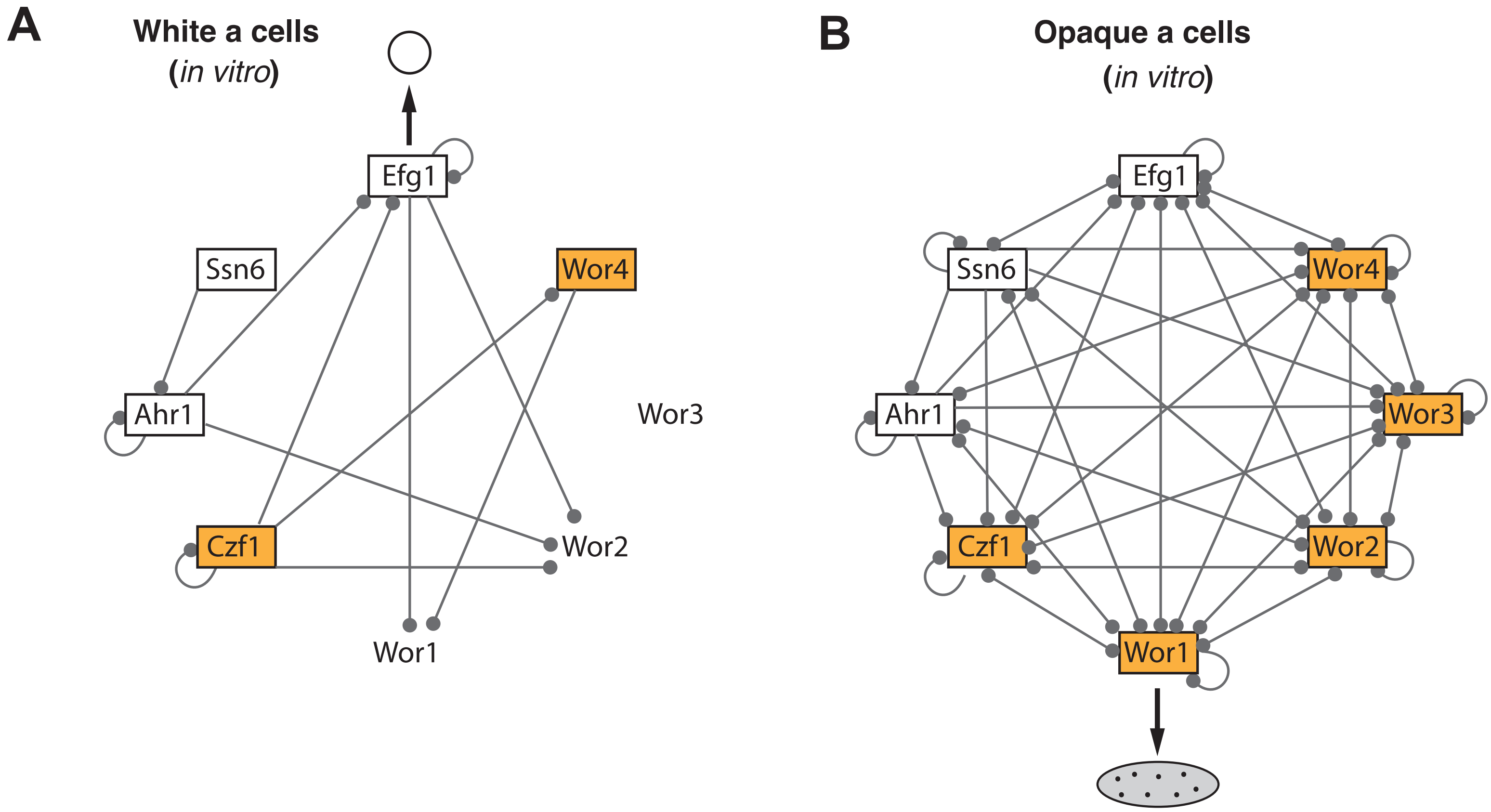

### Figure S2

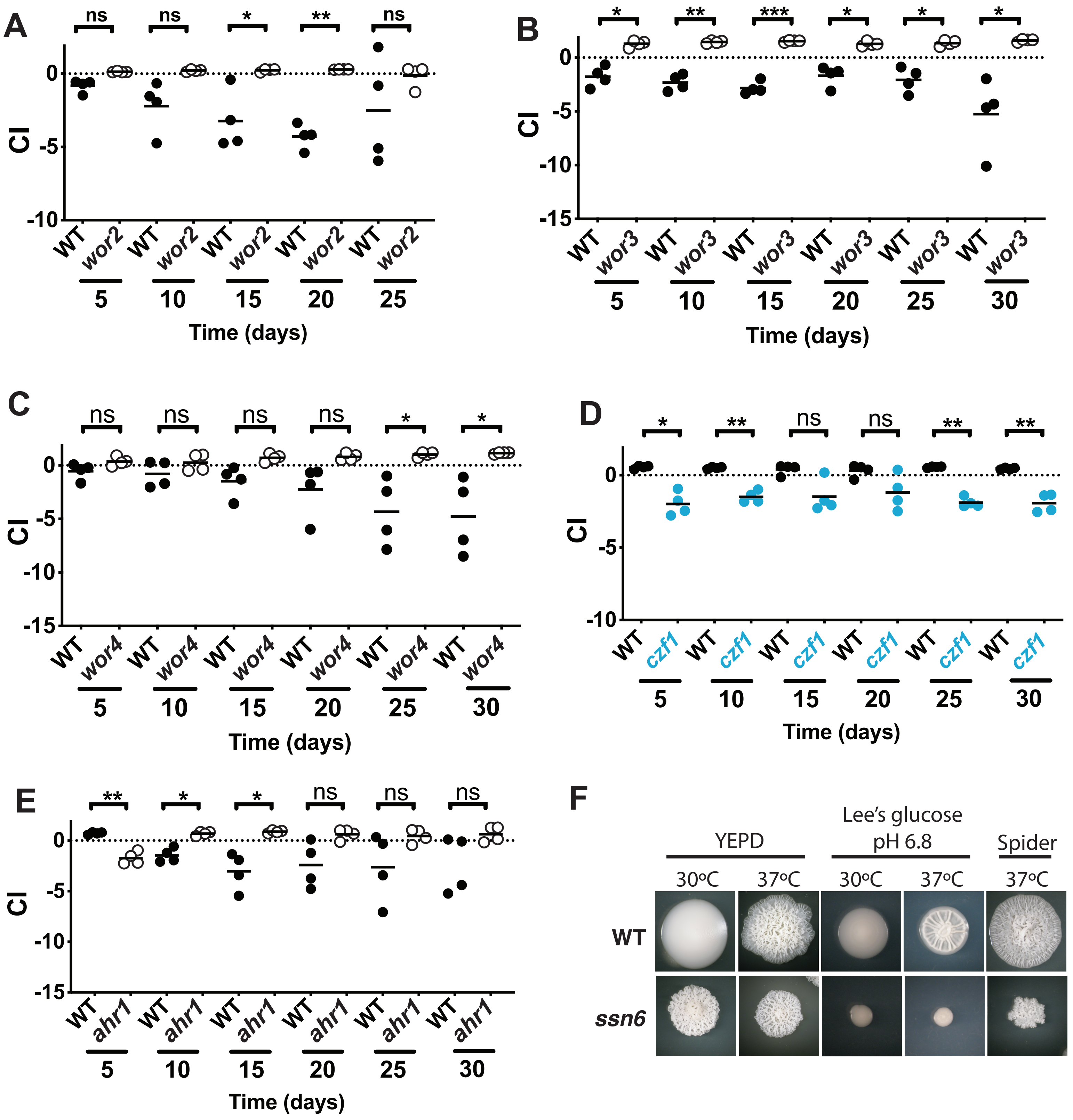

### Figure S3

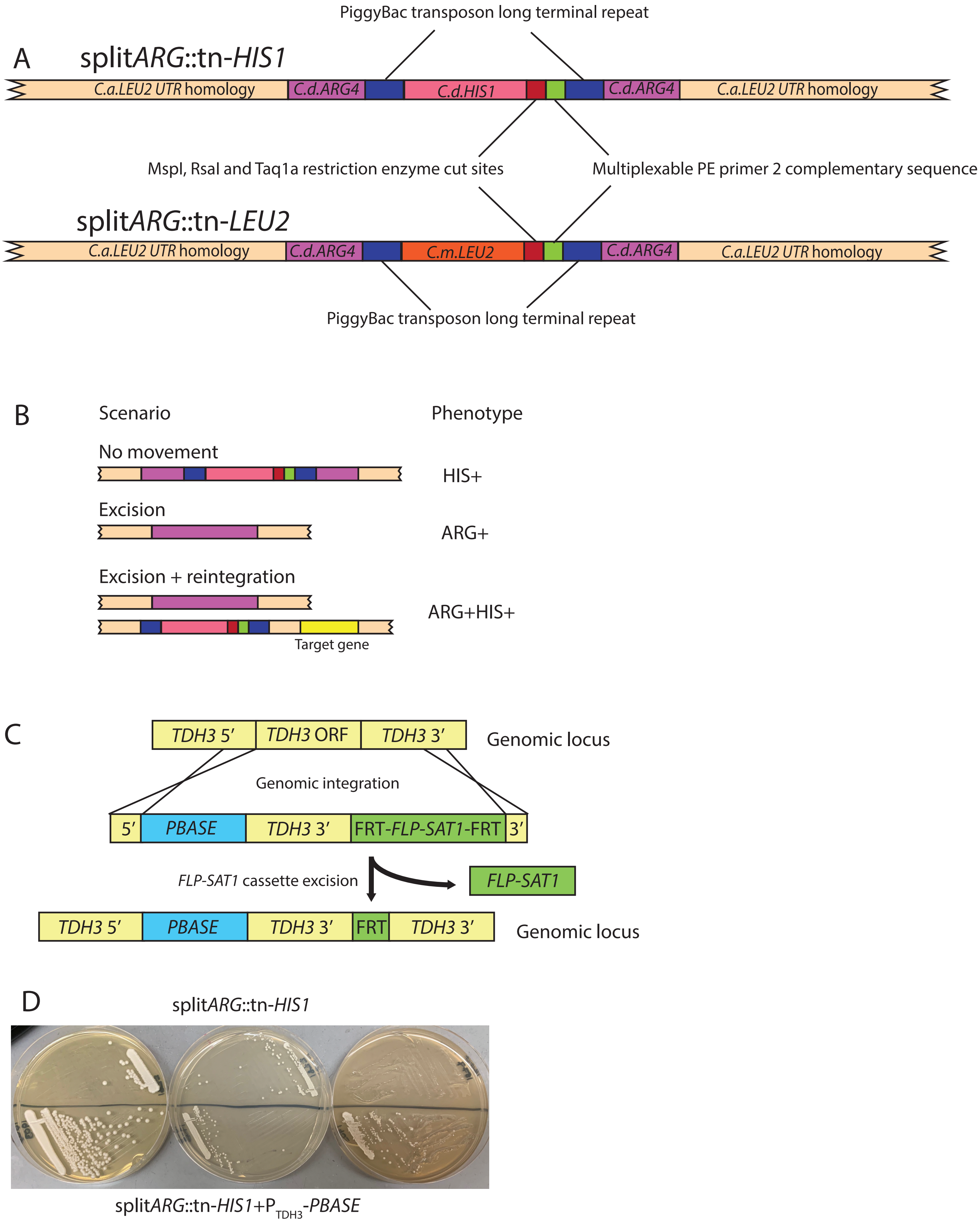
